# Development and characterization of mouse-adapted recombinant SARS-CoV-2 expressing reporter genes

**DOI:** 10.64898/2026.02.04.703885

**Authors:** Sara H. Mahmoud, Nathaniel Jackson, Ramya S. Barre, Yao Ma, Mahmoud Bayoumi, Esteban M. Castro, Shahrzad Ezzatpour, Richard K. Plemper, Stanley Perlman, Chengjin Ye, Luis Martinez-Sobrido

## Abstract

Transgenic K18-hACE2 mice are a standard model for Severe Acute Respiratory Syndrome Coronavirus 2 (SARS-CoV-2), albeit with limitations. A mouse-adapted 30 (MA30) SARS-CoV-2 has been developed to allow infection of wild-type (WT) mice strains. However, SARS-CoV-2 MA30 cannot be easily tracked *in vitro*, *ex vivo*, or *in vivo*. To address the problem, we developed a recombinant (r)SARS-CoV-2 based on the MA30 strain expressing fluorescent (mCherry) and luciferase (nanoluciferase, Nluc) reporter genes, alone or in combination, that enable tracking of viral infection in WT C57BL/6 and BALB/c mice. Insertion of the reporter genes resulted in minor viral attenuation *in vitro*, with ∼0.5-1.0-log lower titers than rSARS-CoV-2 MA30 WT in A549 hACE2 cells, while maintain similar plaque morphology and replication kinetics in Vero AT cells. *In vivo*, reporter-expressing rSARS-CoV-2 MA30 caused transient weight loss, contrasting with lethal rSARS-CoV-2 MA30 WT infection. Bioluminescence imaging of rSARS-CoV-2 MA30 Nluc in C57BL/6 and BALB/c mice revealed peak pulmonary replication at 2 days post-infection, with resolution by day 4, and correlated with tissue viral loads. Our results demonstrate the feasibility of using rSARS-CoV-2 MA30 expressing reporter genes to track viral infection *in vitro, ex vivo*, and *in vivo* without a need for secondary approaches to monitor viral infection as are required for rSARS-CoV-2 MA30 WT. Our system is highly suitable to evaluate prophylactic vaccines and therapeutic antibodies or antiviral approaches in WT or transgenic C57BL/6 and BALB/c mice without the shortcomings of K18-hACE2 mice and with the added advantage of non-invasive monitoring of treatment efficacy.

**Importance:** The K18-hACE2 transgenic mouse model limits the capability to study SARS-CoV-2. While a mouse adapted 30 (MA30) has been developed to study SARS-CoV-2 in wild-type (WT) mice, it does not allow non-invasive tracking of viral infections. Recombinant viruses expressing reporter genes enable real-time monitoring of infection dynamics, opening an avenue to study viral tropism and easily evaluate prophylactic and therapeutic approaches. They furthermore support longitudinal studies, which reduces the number of research animals required. Here, we show that a recombinant (r)SARS-CoV-2 expressing fluorescent (mCherry) and nanoluciferase (Nluc) reporter genes, alone or in combination, can be used to track viral infections *in vitro, ex vivo*, and *in vivo* without the need for secondary approaches that are required to detect SARS-CoV-2 MA30 in WT mice. These reporter-expressing rSARS-CoV-2 MA30 may accelerate vaccine development and antiviral drug discovery in WT or transgenic mice bypassing the need for hACE2 overexpression in K18-hACE2 transgenic mice.

## Introduction

The coronavirus disease 2019 (COVID-19) pandemic, caused by Severe Acute Respiratory Syndrome Coronavirus 2 (SARS-CoV-2), has underscored the critical need for physiologically relevant animal models to study viral pathogenesis, immune responses, and prophylactic and therapeutic interventions (1). Early in the COVID-19 pandemic, transgenic mice expressing human angiotensin-converting enzyme 2 (hACE2), or K18-hACE2 mice, previously developed to study SARS-CoV, became the gold standard for SARS-CoV-2 research due to their susceptibility to viral infection (2, 3). However, the use of K18-hACE2 mice to study SARS-CoV-2 presented some limitations, such as high lethality of SARS-CoV-2 due to overexpression of hACE2 and non-physiological tissue tropism of the human receptor (4). This and other constraints resulted in the development of murine-adapted (MA) SARS-CoV-2 strains capable of infecting wild-type (WT) mice across diverse genetic backgrounds without artificial hACE2 overexpression.

Serial passaging of SARS-CoV-2 in mice resulted in the generation of MA strains with mutations that enhance viral replication. In the case of a SARS-CoV-2 MA10 Washington 1 (WA1) strain, mutations Q493K, Q498H, and N501Y in the Spike (S) glycoprotein have been described to be important for mouse adaptation (5). In the case of the more extensively adapted SARS-CoV-2 MA30 WA1, substitutions K417M, E484K, Q493R, Q498R, and N501Y in the viral S glycoprotein, as well as additional changes in the nonstructural proteins nsp4 (R228E and T295I), nsp8 (S76F), nsp9 (T67A), and ORF8 (S84L) have been identified (**Figure 1A**) (6, 7). These mutations in SARS-CoV-2 MA10 and MA30 improve viral entry and replication in mice, enabling infection of WT BALB/c mice (8). Notably, SARS-CoV-2 MA30 carries key amino acid changes that mirror those observed in human variants of concern (VoC), potentially bridging preclinical findings and clinical outcomes (9). However, current MA SARS-CoV-2 cannot be monitored non-invasively *in vivo* due to the lack of reporter genes that illuminate virus replication, limiting real-time insights into viral dissemination and clearance (6, 7, 10).

**Figure 1.**
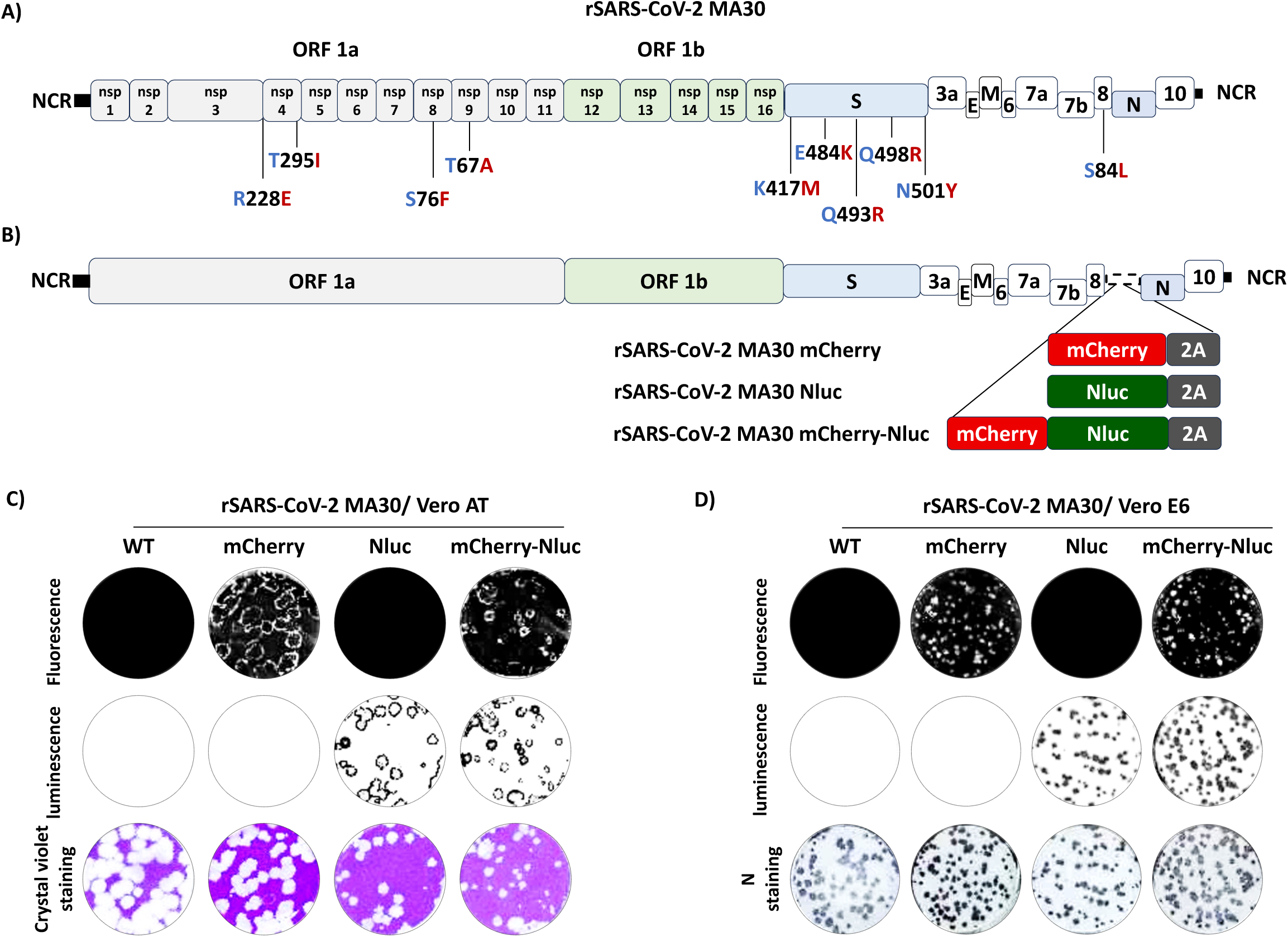
Generation of rSARS-CoV-2 expressing reporter genes and plaque phenotypes. **A)** Amino acid differences between mouse-adapted 30 (MA30) and wild-type (WT) SARS-CoV-2 WA1. Amino acid substitutions identified in SARS-CoV-2 MA30 compared to SARS-CoV-2 WT in the viral open reading frame (ORF) 1a, spike (S), and non-structural protein (nsp8) are indicated; in the mutation labels, the original residue is shown in blue, the position in black, and the substituted residue in red **B)** Schematic representation of rSARS-CoV-2 MA30 expressing mCherry, Nluc, and mCherry-Nluc. Reporter genes were inserted into the 3’ non-coding region (NCR) upstream of the nucleocapsid (N) protein, separated by the PTV-1 2A proteolytic cleavage site. **C-D)** Viral plaques of rSARS-CoV-2 MA30 WT, rSARS-CoV-2 MA30 mCherry, rSARS-CoV-2 MA30 Nluc, and rSARS-CoV-2 MA30 mCherry-Nluc in Vero AT (C) and Vero E6 (D) infected at 3 dpi were observed under a Chemidoc (top), staining with Nluc substrate (middle) or staining with crystal violet (bottom for Vero AT cells), or immunostaining with an antibody against the viral N protein (bottom for Vero E6 cells).

Reporter viruses expressing fluorescent or bioluminescent proteins have revolutionized the study of viral infections by enabling tracking of viral infections *in vitro* and non-invasive, high-resolution visualization of viral replication *in vivo*. For SARS-CoV-2, recombinant viruses expressing different fluorescent (e.g., Venus or mCherry) and nanoluciferase (Nluc) proteins have been previously described (11). Initially, rSARS-CoV-2 reporter strains were generated by introducing the fluorescent or Nluc reporter genes at the position of the viral open reading frame (ORF)7a protein (12–15). However, we have described rSARS-CoV-2 expressing the same reporter genes as a fusion with the viral nucleocapsid (N) protein, separated by the porcine teschovirus 1 (PTV-1) 2A proteolytic cleavage site (9). This approach allowed higher expression of the reporter genes without affecting viral replication or the need for substituting any viral protein with the reporter gene (16). Moreover, we have demonstrated the feasibility of generating an rSARS-CoV-2 dual reporter strain that expresses both fluorescent (mCherry) and luciferase (Nluc) as a fusion protein from the locus of the viral N gene, using the same PTV-1 2A approach (17). These reporter rSARS-CoV-2 have been recognized as an excellent tool to track viral infections *in vitro, ex vivo*, and *in vivo* (18). However, the use of these reporter-expressing rSARS-CoV-2 based on the original WA-1 strain was limited to K18 hACE2 transgenic mice (19).

Here, we addressed the problem by generating rSARS-CoV-2 MA30s expressing mCherry, Nluc, or mCherry-Nluc, which we characterized *in vitro, ex vivo*, and *in vivo* in C57BL/6 and BALB/c mice. *In vitro*, the rSARS-CoV-2 MA30 expressing reporter genes retained viral replication and plaque phenotype similar to that of rSARS-CoV-2 MA30 WT. *In vivo*, the rSARS-CoV-2 MA30 expressing reporter genes allowed viral infection in C57BL/6 and BALB/c mice, circumventing the need for hACE2 overexpression in K18-hACE2 transgenic mice, as is required for parental strains of SARS-CoV-2 (20). Importantly, the recombinant viruses expressing Nluc (rSARS-CoV-2 Nluc and rSARS-CoV-2 mCherry-Nluc) allowed us to monitor viral infection in intact mice using an *in vivo* imaging system (IVIS). We also tracked viral infection *ex vivo* in the lungs of C57BL/6 and BALB/c mice infected with rSARS-CoV-2 Nluc, rSARS-CoV-2 mCherry, or rSARS-CoV-2 mCherry-Nluc through IVIS. However, rSARS-CoV-2 MA30 expressing reporter genes were attenuated compared to rSARS-CoV-2 MA30 WT, resulting in lower pathogenicity, and both C57BL/6 and BALB/c mice infected with rSARS-CoV-2 MA30 expressing reporter genes survived the infections.

By combining murine adaptations present in SARS-CoV-2 MA30 with fluorescent and/or luciferase reporter expression and our bacterial artificial chromosome (BAC)-based reverse genetics approaches to generate rSARS-CoV-2, we have generated rSARS-CoV-2 MA30s expressing mCherry and/or Nluc proteins that facilitate monitoring viral infection *in vivo* and *ex vivo* in WT C57BL/6 and BALB/c mice without the need for K18-hACE2 transgenic mice. This work supports the evaluation of viral pathogenicity in real time and advances the discovery of new prophylactics and therapeutics for the treatment of SARS-CoV-2 infection.

## Results

### Rescue of WT and reporter-expressing rSARS-CoV-2 MA30

Serial passaging of SARS-CoV-2 WA1 in BALB/c mice resulted in a mouse-adapted strain, MA30 (21, 22). Sequence analysis of SARS-CoV-2 MA30 revealed mutations in the S (K417M, E484K, Q493R, Q498R, and N501Y), nsp4 (R228E and T295I), nsp8 (S76F), nsp9 (T67A), and ORF8 (S84L) proteins (**Figure 1A**) (23–25). We and others have shown that recombinant viruses, including SARS-CoV-2, expressing reporter fluorescent and/or luciferase proteins, represent an excellent tool to track infections *in vitro, ex vivo*, and *in vivo* (11, 26–28). Thus, we employed reverse genetics to develop rSARS-CoV-2 MA30 expressing fluorescent (mCherry) and bioluminescent (Nluc) reporter proteins, alone or in combination, to facilitate the study of SARS-CoV-2 MA30 infection in parental murine models (28) (**Figure 1B**). The reporter genes were inserted upstream of the nucleocapsid (N) protein, separated by the porcine teschovirus-1 (PTV-1) 2A proteolytic sequence to ensure the independent expression of N and reporter proteins (**Figure 1B**). We have previously used this approach to generate rSARS-CoV-2 WA1 expressing fluorescent and Nluc reporter genes, likewise alone or in combination (9, 16, 28, 29). We transfected the corresponding BACs into Vero E6 cells to generate the rSARS-CoV-2 MA30 mCherry, rSARS-CoV-2 MA30 Nluc, and rSARS-CoV-2 MA30 Nluc-mCherry variants. As reference for our studies, we also generate a rSARS-CoV-2 MA30 WT.

### *In vitro* characterization of WT and reporter-expressing rSARS-CoV-2 MA30

To characterize the *in vitro* properties of the reporter-expressing rSARS-CoV-2 MA30 viruses, we first examined their plaque phenotypes in Vero AT and Vero E6 cells **(Figures 1C and 1D**, respectively). All reporter-expressing rSARS-CoV-2 MA30 produced visible plaques in both cell lines at 3 days post-infection (dpi). Notably, the plaque size was comparable, although slightly smaller, to that of rSARS-CoV-2 MA30 WT (**Figures 1C and 1D**). As expected, we also observed differences in plaque size and morphology between the two cell lines. In Vero AT cells, viral plaques were larger and more clearly defined (**Figure 1C**). In contrast, viral plaques in Vero E6 cells appeared smaller and less distinct (**Figure 1D**). This difference in plaque morphology suggests that the viruses spread more efficiently in Vero AT cells compared to Vero E6 cells. We have observed similar results with different natural SARS-CoV-2 strains that replicate more efficiently and produce larger plaques in Vero AT cells compared to Vero E6 cells (30). Importantly, within each cell line, the plaque morphology was consistent across all four viruses, indicating that the insertion of the reporter gene(s) did not significantly alter viral fitness or cell-to-cell spread, as previously described for SARS-CoV-2 WA1 (17). To verify the genetic integrity of the reporter gene inserts, RT-PCR analysis was performed on viral RNA extracted from individual plaques isolated from rSARS-CoV-2 MA30 mCherry, rSARS-CoV-2 MA30 Nluc, and rSARS-CoV-2 MA30 mCherry-Nluc infected cells at passage 1 (P1). Specific primers targeting the flanking regions of the reporter genes were used to amplify the inserted sequences via RT-PCR. PCR products were sequenced to confirm the integrity of the inserts and the correct positioning upstream of the N protein. All examined plaque isolates yielded PCR products of the expected size and sequence identity, confirming the correct insertion of the reporter genes in the viral genome (data not shown). These results are consistent with previous studies using the same PTV-1 2A cleavage approach with SARS-CoV-2 WA1, which demonstrated the insertion and stability of the reporter gene constructs (12, 29).

We next assessed viral growth kinetics of WT and reporter-expressing rSARS-CoV-2 MA30 in Vero AT and A549 hACE2 cells (**Figures 2A–2F**). In Vero AT cells, rSARS-CoV-2 MA30 WT replicated more efficiently and reached significantly higher titers than reporter-expressing rSARS-CoV-2 MA30 at all time points (12, 24, 48, and 72 hpi), peaking at approximately 10 PFU/mL by 48 hpi compared to 10^5.6^-10^5.9^ PFU/mL for reporter-expressing rSARS-CoV-2 MA30 (**Figure 2A**). At 24 hpi, rSARS-CoV-2 MA30 mCherry exceeded titers of rSARS-CoV-2 MA30 Nluc and rSARS-CoV-2 MA30 mCherry-Nluc, while at 72 hpi, rSARS-CoV-2 MA30 mCherry showed lower titers than rSARS-CoV-2 MA30 Nluc and rSARS-CoV-2 MA30 mCherry-Nluc (**Figure 2A**). In contrast, in A549 hACE2 cells, viral titers were comparable at 12 hpi across all strains (∼10^2.3^ PFU/mL). Significant differences were observed only at 24 hpi, when rSARS-CoV-2 MA30 WT and rSARS-CoV-2 M30 mCherry (∼10^3^ PFU/mL) exceeded viral titers of rSARS-CoV-2 MA30 Nluc and rSARS-CoV-2 MA30 mCherry-Nluc (**Figure 2B**). By 48 and 72 hpi, all viruses reached similar titers with no significant differences (**Figure 2B**). These results revealed cell line-dependent replication kinetics: in Vero AT cells, rSARS-CoV-2 MA30 WT reached higher titers than all the reporter-expressing rSARS-CoV-2 MA30 across all time points, whereas in A549 hACE2 cells, reporter-expressing rSARS-CoV-2 MA30 replication was comparable to rSARS-CoV-2 WT after 24 hpi, suggesting that the fitness penalty associated with reporter gene expression was transient and cell type dependent. To validate Nluc expression and assess viral replication dynamics, we measured Nluc activity in cell culture supernatants from infected Vero AT and A549 hACE2 cells (**Figures 2C and 2D**, respectively). Only rSARS-CoV-2 MA30 Nluc and rSARS-CoV-2 MA30 mCherry-Nluc produced robust Nluc signals at all time points, with no detection above background in rSARS-CoV-2 MA30 WT or rSARS-CoV-2 MA30 mCherry infected cell culture supernatants in Vero AT (**Figure 2C**) or A549 hACE2 (**Figure 2D**). Notably, Nluc activity from cells infected with rSARS-CoV-2 MA30 mCherry-Nluc was consistently higher than from cells infected with rSARS-CoV-2 MA30 Nluc (**Figures 2C and 2D**). Importantly, Nluc kinetics in both cell lines showed similar temporal profiles despite marked differences in viral titers and mCherry expression dynamics.

**Figure 2.**
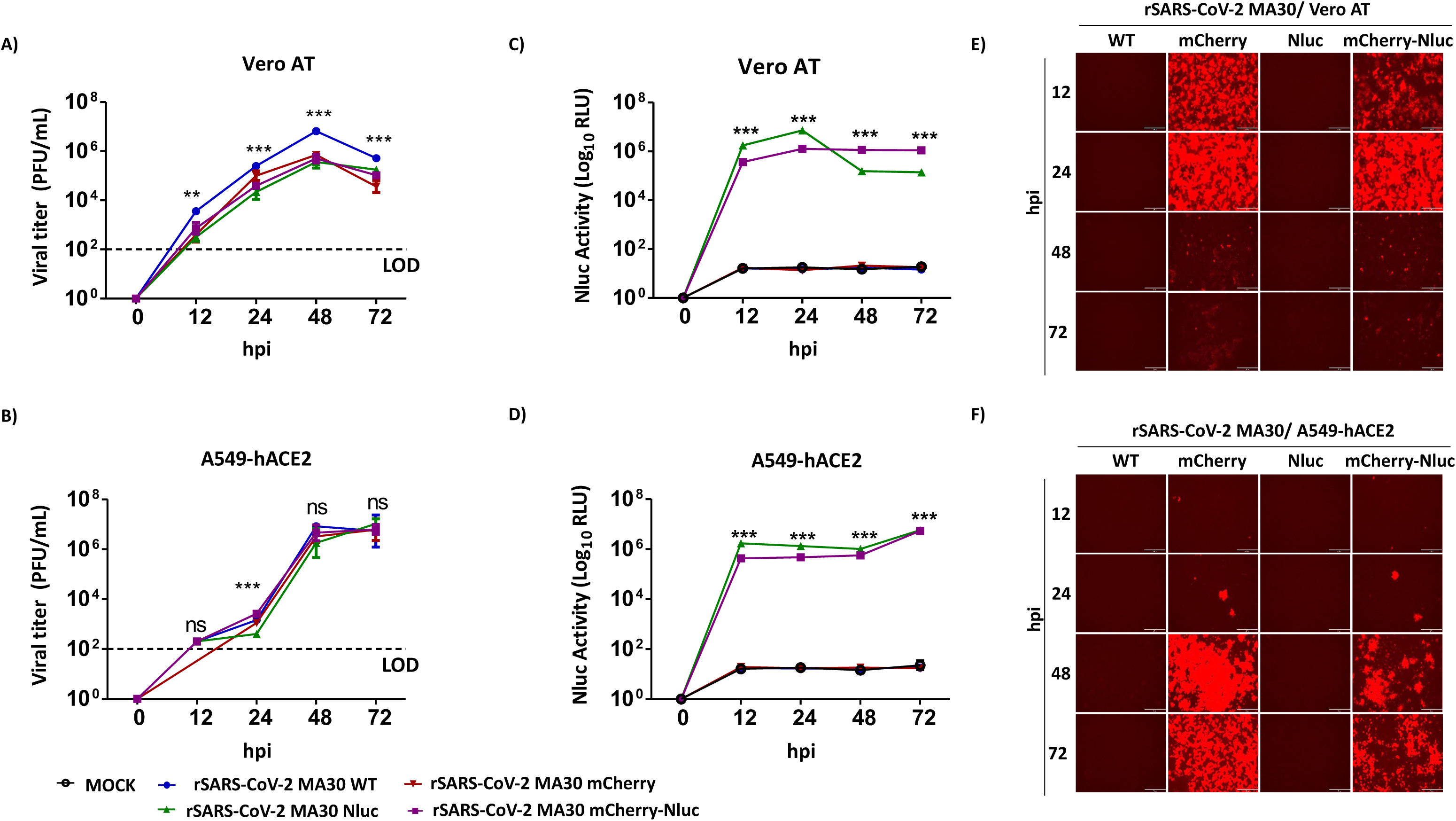
*In vitro* characterization of rSARS-CoV-2 MA30 expressing reporter genes. **A-B):** Viral titers (PFU/mL) in the cell culture supernatants of Vero AT (A) and A549 hACE2 (B) cells infected (MOI, 0.01) with rSARS-CoV-2 MA30 WT, rSARS-CoV-2 MA30 mCherry, rSARS-CoV-2 MA30 Nluc, and rSARS-CoV-2 MA30 mCherry-Nluc at the indicated hours post-infection (hpi) were determined by plaque assay. Data represent the mean values and SD of triplicates. LOD, limit of detection. *\*\*p* < 0.01; *\*\*\*p* < 0.001; ns, not significant. **C-D)** Nluc expression in the same cell culture supernatants obtained from Vero AT (A) and A549 hACE2 (B) infected cells is represented in relative light units (RLU). *\*\*\*p* < 0.001. **E-F)** Vero AT **(E)** and A549 hACE2 **(F)** cells were infected (MOI, 0.01) with rSARS-CoV-2 MA30 WT, rSARS-CoV-2 MA30 mCherry, rSARS-CoV-2 MA30 Nluc, or rSARS-CoV-2 MA30 mCherry-Nluc. At the indicated times post-infection, mCherry expression was visualized under a fluorescence microscope. Representative images are shown. Scale bars = 300 µm. Magnification = 10X.

We also examined mCherry expression kinetics in Vero AT (**Figure 2E**) and A549 hACE2 (**Figure 2F**) cells infected with the different rSARS-CoV-2 MA30 viruses. In Vero AT cells, both rSARS-CoV-2 MA30 mCherry and rSARS-CoV-2 MA30 mCherry-Nluc became mCherry fluorescence-positive as early as 12 hpi, with signal intensity increasing at 24 hpi and declining by 48 hpi. In contrast, mCherry expression in A549 hACE2 cells infected with rSARS-CoV-2 MA30 mCherry and rSARS-CoV-2 MA30 mCherry-Nluc was first detected at 24 hpi and became more pronounced at 48 and 72 hpi. This cell-type-dependent difference in mCherry expression kinetics directly reflected the cell-specific differences in viral dynamics (**Figures 2A and 2B**): the earlier and more rapid mCherry signal in Vero AT cells (**Figure 2E**) paralleled the higher and earlier viral replication observed in this cell line (**Figure 2A**) compared to mCherry expression (**Figure 2F**) and viral titers (**Figure 2B**) observed in A549 hACE2 cells. This finding was contrary to Nluc expression that always reached comparable values in Vero AT (**Figure 2C**) and A549 HACE2 (**Figure 2D**) post-infection.

These results indicate that the reporter-expressing rSARS-CoV-2 MA30 maintained efficient replication in Vero AT and A549 hACE2 cells, with only minor attenuation compared to rSARS-CoV-2 MA30 WT. Importantly, reporter gene (mCherry and/or Nluc) expression allowed tracking and detection of viral infection using fluorescent microscopy (mCherry) or fluorescent plate readers (Nluc), providing qualitative and quantitative measurements of viral infection and replication without a need for secondary approaches to detect the presence of rSARS-CoV-2 in infected cells, making these reporter-expressing virus strains valuable tools for *in vitro* studies.

### Age- and strain-dependent susceptibility of rSARS-CoV-2 MA30 in mice

We next compared three mouse strains (C57BL/6, BALB/c, and K18-hACE2) at two age groups (6 and 21 weeks-old) using three infection doses (10^2^, 10^3^, and 10^4^ PFU/mouse) of rSARS-CoV-2 MA30 WT (**Figure 3**). Younger mice (6 weeks-old) (**Figures 3A-3C**) consistently showed lower susceptibility to rSARS-CoV-2 MA30 WT infection than older mice (21 weeks-old) (**Figures 3D-3F**) across all mouse strains and viral doses, as evidenced by less severe weight loss and higher survival rates. This age-dependent effect was particularly pronounced in C57BL/6 **(Figures 3A and 3D**), followed by BALB/c (**Figures 3B and 3E**) mice. Among the different mouse strains tested, K18-hACE2 mice exhibited the highest susceptibility to rSARS-CoV-2 MA30 WT. Infection resulted in rapid weight loss and 100% mortality observed at all doses in both age groups, except in 6-week-old K18 hACE2 mice infected with the lower doses of 10^3^ and 10^2^ PFU, which showed 20% and 100% survival, respectively (**Figures 3C and 3F**).

**Figure 3.**
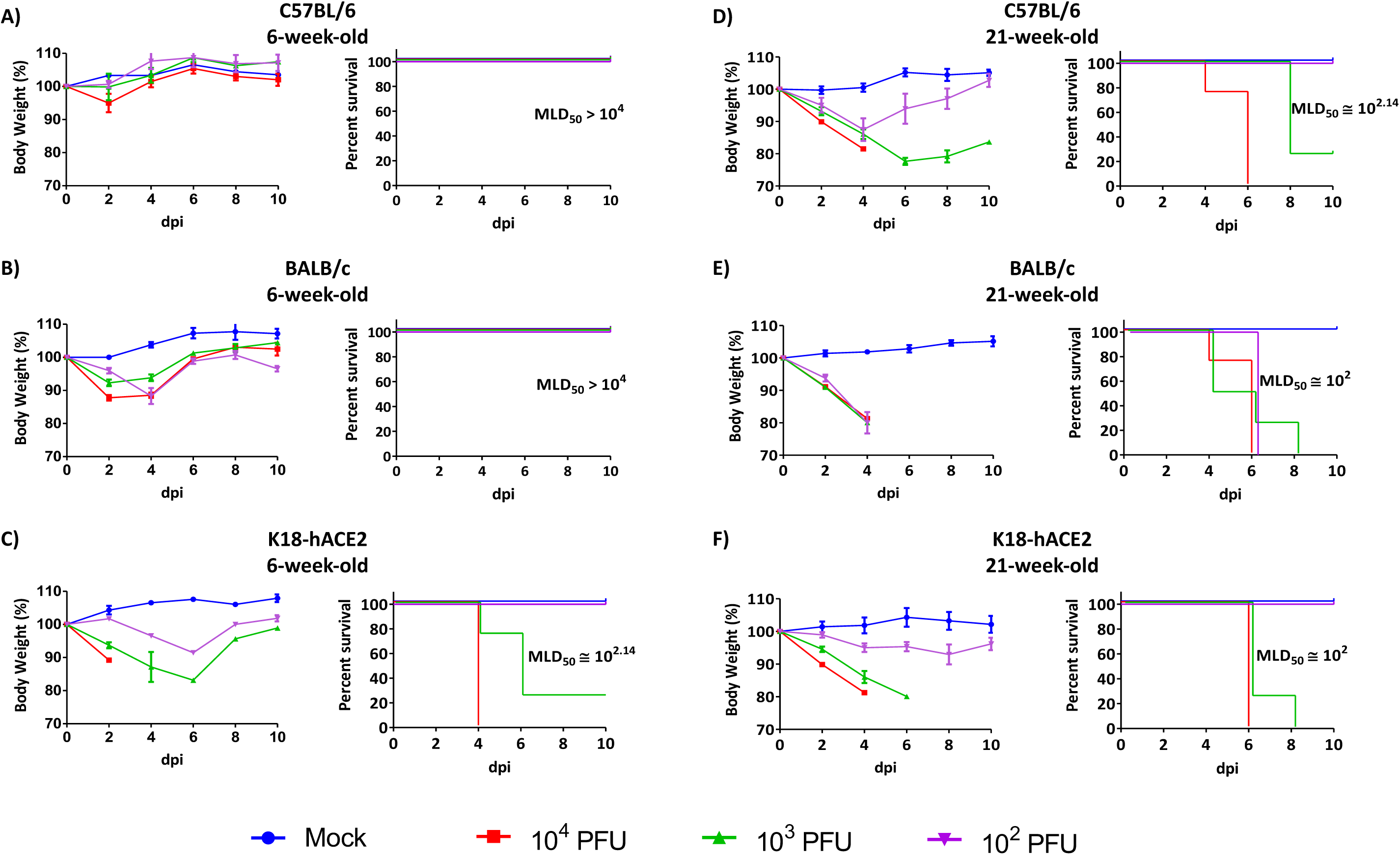
Age-dependent and strain-specific mouse susceptibility to rSARS-CoV-2 MA30 WT infection. **A-C)** C57BL/6 **(A),** BALB/c **(B),** and K18-hACE2 **(C)** 6-week-s old mice were infected intranasally (i.n.) with 10^2^, 10^3^, or 10^4^ PFU of rSARS-CoV-2 MA30 WT. Body weight changes (left panel) and survival (right panel) were monitored for 10 days after viral infection. **D-F)** C57BL/6 **(D),** BALB/c **(E),** and K18-hACE2 **(F)** 21-week-old mice were infected with 10^2^, 10^3^, or 10^4^ PFU of rSARS-CoV-2 MA30 WT. Body weight changes (left panel) and survival rates (right panel) were monitored over 10 days. Data represent mean ± SD (n = 4 mice/group).

C57BL/6 mice displayed the highest resistance to rSARS-CoV-2 MA30 WT infection in both age groups, with minimal weight loss and 100% survival in young mice at all viral doses (**Figure 3A**). BALB/c mice demonstrated intermediate susceptibility, showing more severe outcomes than C57BL/6 mice, especially at higher doses and in older animals (**Figures 3B and 3E**). We observed a clear viral inoculum-dependent effect on disease severity. Infection with the higher viral amount (10^4^ PFU) resulted in severe weight loss and lower survival rates across all mouse strains in the aged group (**Figures 3D, 3E, and 3F**). This inoculum effect was most prominent in BALB/c mice, where increasing amounts led to progressively worse outcomes. Based on these results, we were able to determine the median lethal inoculum (MLD_50_) of rSARS-CoV-2 MA30 WT in C57BL/6, BALB/c, and K18-hACE2 mice at 6 and 21 weeks of age. In 6-week-old C57BL/6 and BALB/c mice, rSARS-CoV-2 MA30 WT exhibited an MLD_50_ > 10^4^ PFU (**Figures 3A and 3B**). Conversely, in 6-week-old K18-hACE2 mice, rSARS-CoV-2 MA30 WT exhibited an MLD_50_ of approximately 10^2.14^ PFU (**Figure 3C**), comparable to that observed with a natural, non-mouse-adapted, SARS-CoV-2 WA1 strain in K18-hACE2 mice (31–34). In 21-week-old C57BL/6 mice, rSARS-CoV-2 MA30 WT showed_₅₀_ of approximately 10 PFU (**Figure 3D**). A similar MLD_₅₀_ (∼10 PFU) was observed in age-matched BALB/c mice (**Figure 3E**). In 21-week-old K18-hACE2 mice, the MLD was also approximately 10^2^ PFU (**Figure 3F**), comparable to that previously observed in 6-week-old K18-hACE2 animals. When considered across strains and age groups, K18-hACE2 mice displayed high susceptibility regardless of age. In contrast, both C57BL/6 and BALB/c mice showed increased susceptibility at 21 weeks compared with younger animals, and BALB/c mice were consistently more susceptible than C57BL/6. Based on these observations, 21-week-old BALB/c and C57BL/6 mice were selected as a suitable model for subsequent pathogenicity studies using WT or reporter-expressing rSARS-CoV-2 MA30.

### Characterization of reporter-expressing rSARS-CoV-2 MA30 in C57BL/6 mice

To investigate the *in vivo* kinetics of reporter-expressing rSARS-CoV-2 MA30, 22-week-old female C57BL/6 (n = 5) were infected i.n. with 10^5^ PFU/mouse of rSARS-CoV-2 MA30 mCherry, rSARS-CoV-2 MA30 Nluc, and rSARS-CoV-2 MA30 mCherry-Nluc. Mock-infected mice and mice infected with rSARS-CoV-2 MA30 WT were included as controls. Then, Nluc activity was evaluated in intact animals at 1, 2, 4, and 6 dpi with an Ami HT IVIS (**Figure 4A**). The same mice were monitored for changes in body weight (**Figure 4B**) and survival (**Figure 4C**) for 10 days after viral infection. *In vivo* imaging revealed Nluc activity primarily in the lungs at 2 dpi in mice infected with rSARS-CoV-2 MA30 Nluc and rSARS-CoV-2 MA30 mCherry-Nluc, with Nluc signal decreasing by 4 dpi (**Figure 4A**). Mice infected with rSARS-CoV-2 MA30 WT showed the most severe weight loss, reaching approximately 75% of initial body weight by 5 dpi, resembling our previous results (**Figure 3D**). Interestingly, mice infected with the reporter-expressing rSARS-CoV-2 MA30 exhibited less severe weight loss compared to those infected with rSARS-CoV-2 MA30 WT, and all infected animals recovered after 4-5 dpi. Mock-infected mice maintained stable body weights throughout the entire experiment. Survival data (**Figure 4C**) indicated 100% mortality in C57BL/6 mice infected with rSARS-CoV-2 MA30 WT by 6 dpi, similar to our previous study (**Figure 3D**). In contrast, all C57BL/6 mice infected with the reporter-expressing rSARS-CoV-2 MA30 survived the infection, as did the mock-infected controls. Notably, all the C57BL/6 mice infected with the rSARS-CoV-2 MA30 expressing reporter-genes loss ∼10% of their initial body weight. These results suggest that the reporter-expressing rSARS-CoV-2 MA30 were attenuated compared to rSARS-CoV-2 MA30 WT in C57BL/6 mice. However, contrary to rSARS-CoV-2 MA30 WT, both Nluc-expressing rSARS-CoV-2 MA30 provide a powerful tool for non-invasive monitoring of viral infection kinetics in live animals, offering valuable insights into the spatiotemporal dynamics of SARS-CoV-2 infection in the C57BL/6 mouse model.

**Figure 4.**
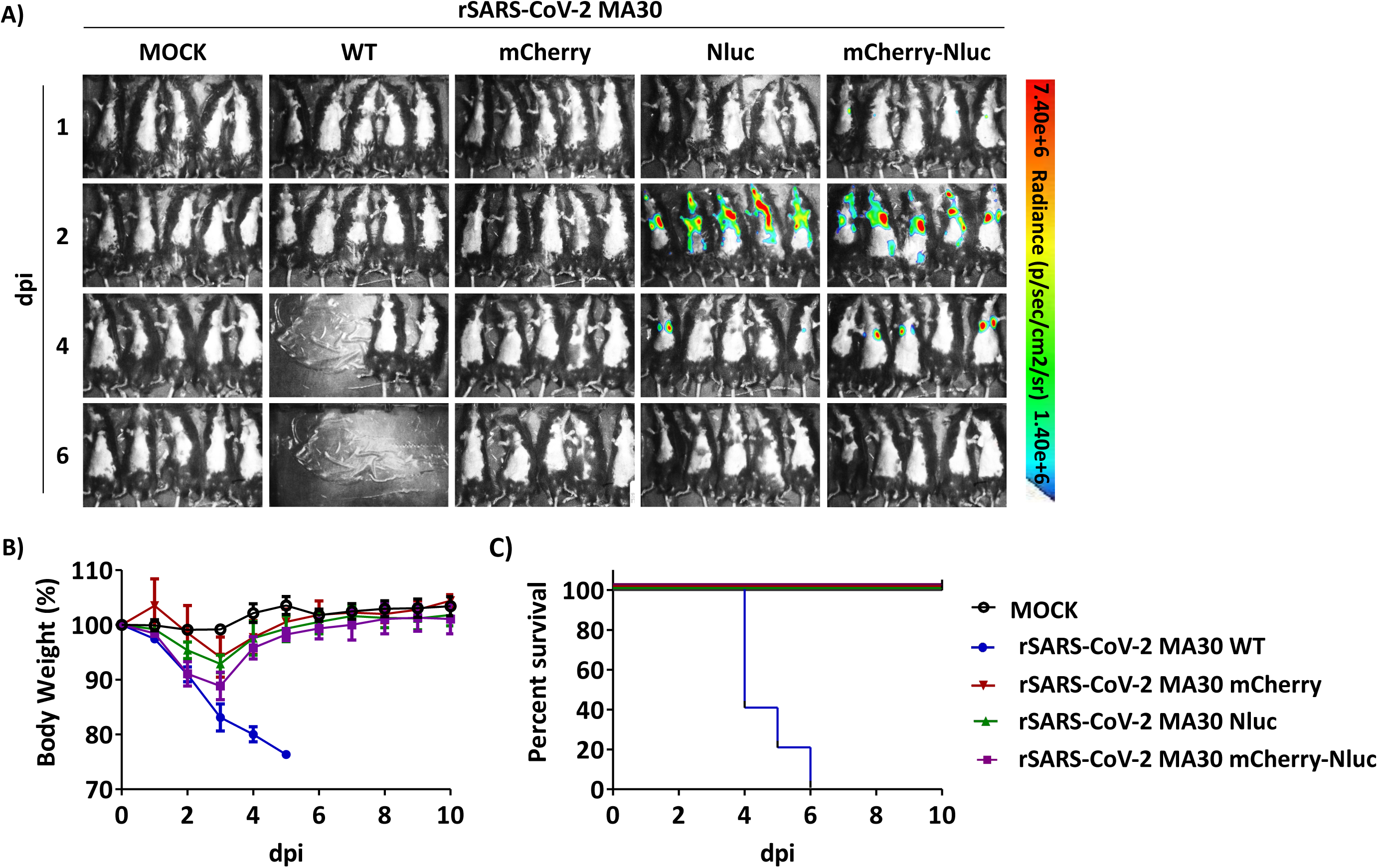
*In vivo* kinetics of rSARS-CoV-2 MA30 reporter viruses in C57BL/6 mice: **A)** C57BL/6 mice 22-week-old (n=5/group) were mock-infected or infected i.n. with 10^5^ PFU/mouse of rSARS-CoV-2 MA30 WT, rSARS-CoV-2 MA30 mCherry, rSARS-CoV-2 MA30 Nluc, or rSARS-CoV-2 MA30 mCherry-Nluc. Nluc expression in the whole mouse at the indicated dpi was evaluated with an Ami HT IVIS. Representative images of the same mouse at 1, 2, 4, and 6 dpi are shown. **B-C)** Body weight loss **(B)** and survival **(C)** of mice infected in panel A were monitored for 10 days after viral infection.

In a separate experiment, we further followed Nluc signals in C57BL/6 mice infected with rSARS-CoV-2 MA30 Nluc and rSARS-CoV-2 MA30 Nluc-mCherry (**Figure 5**). *In vivo* bioluminescence imaging (**Figure 5A**) revealed strong Nluc activity in animals infected with rSARS-CoV-2 MA30 Nluc and rSARS-CoV-2 MA30 mCherry-Nluc at 2 dpi, primarily localized in the lungs. Signals decreased by 4 dpi, especially in C57BL/6 mice infected with rSARS-CoV-2 MA30 Nluc (**Figure 5C**). As expected, no bioluminescence was observed in mock-infected mice or animals infected with rSARS-CoV-2 MA30 WT or rSARS-CoV-2 MA30 mCherry at either 2 or 4 dpi (**Figures 5A and 5C**). *Ex vivo* imaging of excised lungs from the same animal groups at 2 and 4 dpi (**Figures 5B and 5D**, respectively) confirmed the presence of Nluc in lungs from C57BL/6 mice infected with Nluc-expressing rSARS-CoV-2 MA30. Signal intensity correlated well with *in vivo* imaging of intact mice (**Figures 5A and 5C**). Additionally, fluorescence imaging of lungs from mice infected with rSARS-CoV-2 MA30 mCherry and rSARS-CoV-2 MA30 mCherry-Nluc exhibited distinct mCherry expression by 2 dpi, demonstrating expression of the fluorescent protein in the infected animals. Comparable to mice infected with rSARS-CoV-2 MA30 Nluc, we did not detect mCherry expression in C57BL/6 mice infected with rSARS-CoV-2 MA30 mCherry or rSARS-CoV-2 MA30 mCherry-Nluc at 4 dpi. No mCherry signal was observed in the lungs of mock-infected C57BL/6 mice or in the lungs of C57BL/6 mice infected with rSARS-CoV-2 MA30 WT or rSARS-CoV-2 MA30 Nluc.

**Figure 5.**
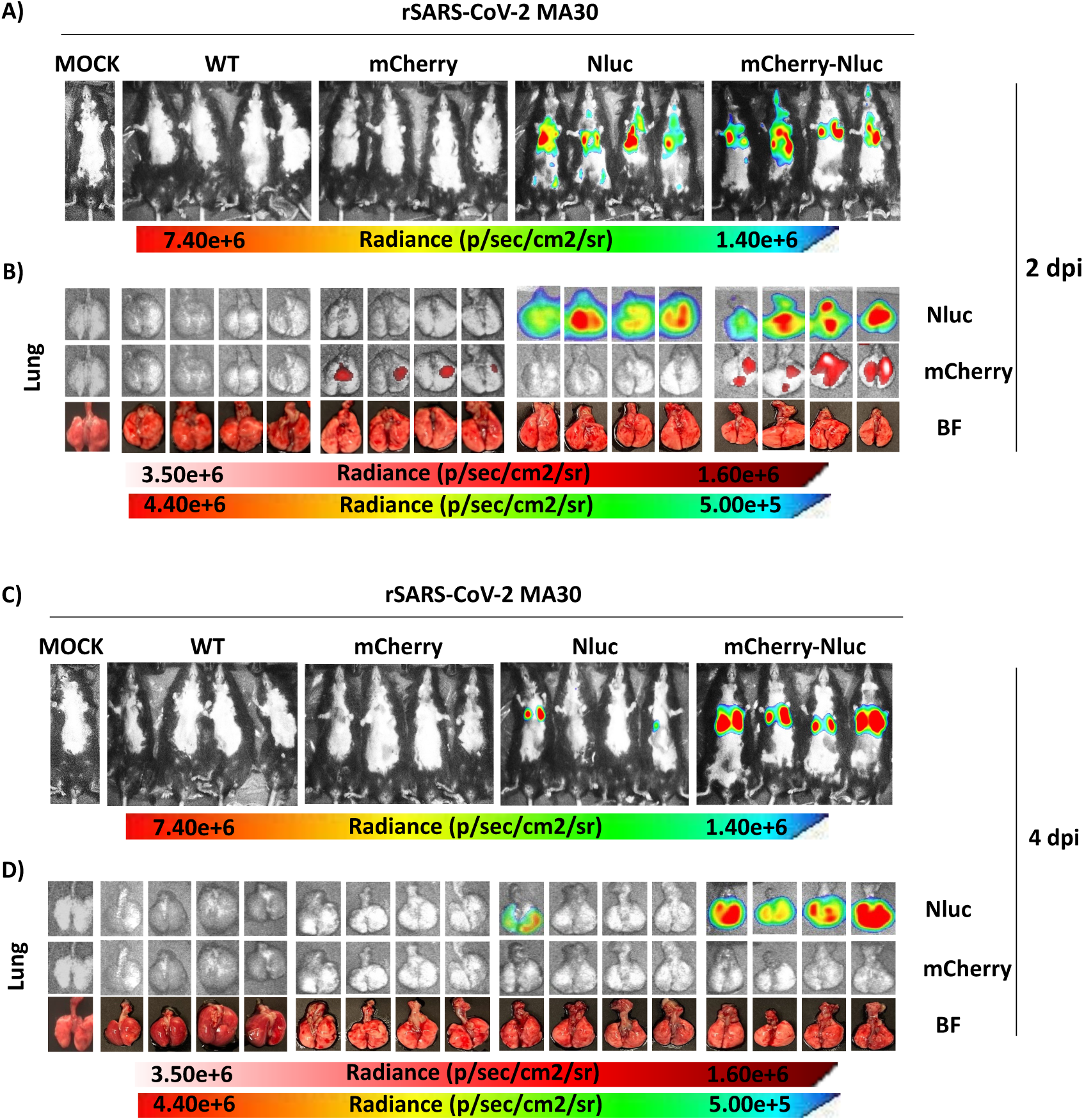
*In vivo* and *ex vivo* kinetics of reporter-expressing rSARS-CoV-2 MA30 in C57BL/6 mice: **A and C)** Nluc activity in whole 22-week-old C57BL/6 mice mock-infected or infected i.n. with 10^5^ PFU/mouse with rSARS-CoV-2 MA30 WT, rSARS-CoV-2 MA30 mCherry, rSARS-CoV-2 MA30 Nluc, and rSARS-CoV-2 MA30 mCherry-Nluc at 2 (n=4 mice/group) and 4 (n=4 mice/group) dpi using the Ami HT IVIS. **B and D)** Excised lungs from mock-infected or infected mice from panels A and C were monitored for Nluc and mCherry expression at 2 and 4 dpi. Bright field (BF) images of the lungs were also collected at the same time points.

Viral titers in lung and nasal turbinate tissue homogenates from the same infected C57BL/6 mice correlated with *in vivo* Nluc and *ex vivo* Nluc and mCherry expression (**Figure 6A**). At 2 dpi, lung viral titers were higher in rSARS-CoV-2 MA30 WT-infected mice (∼10^8^ PFU/mL), followed by reporter-expressing rSARS-CoV-2 MA30 (∼10^7^ PFU/mL). By 4 dpi, titers decreased in all animal groups, and rSARS-CoV-2 MA30 expressing reporter genes showed a more pronounced reduction in viral titers (∼10^3^ PFU/mL) than those observed with rSARS-CoV-2 MA30 WT (∼10^6^ PFU/mL). Viral titers in the nasal turbinate followed a similar pattern, with higher viral titers in C57BL/6 mice infected with rSARS-CoV-2 MA30 WT at both 2 and 4 dpi than in mice infected with the reporter-expressing rSARS-CoV-2 MA30 (**Figure 6A**). Viral titers in the brain of infected mice were low, but detectable, in C57BL/6 mice infected with rSARS-CoV-2 MA30 WT at both 2 and 4 dpi (**Figure 6A**). Only viral titers in the brain of two C57BL/6 mice infected with rSARS-CoV-2 MA30 Nluc were detected at 2 dpi (**Figure 6A**). Quantification of Nluc activity in the same tissue homogenates (**Figure 6B**) showed the highest levels of expression in lungs and nasal turbinate of C57BL/6 mice infected with rSARS-CoV-2 MA30 Nluc and rSARS-CoV-2 MA30 mCherry-Nluc. Lung Nluc activity peaked at 2 dpi and decreased by 4 dpi, similar to viral titers (**Figure 6A**), while activity in the nasal turbinate remained relatively stable, although declined by 4 dpi (**Figure 6B**). Low, but detectable, Nluc activity was observed in brain homogenates of C57BL/6 mice infected with rSARS-CoV-2 MA30 Nluc and rSARS-CoV-2 MA30 mCherry-Nluc at 2 and 4 dpi, suggesting potential infection of the brain (**Figure 6B**). As expected, we did not detect Nluc expression in any of the tissue homogenates of C57BL/6 mice infected with rSARS-CoV-2 MA30 WT, rSARS-CoV-2 MA30 mCherry, or in mock-infected animals.

**Figure 6.**
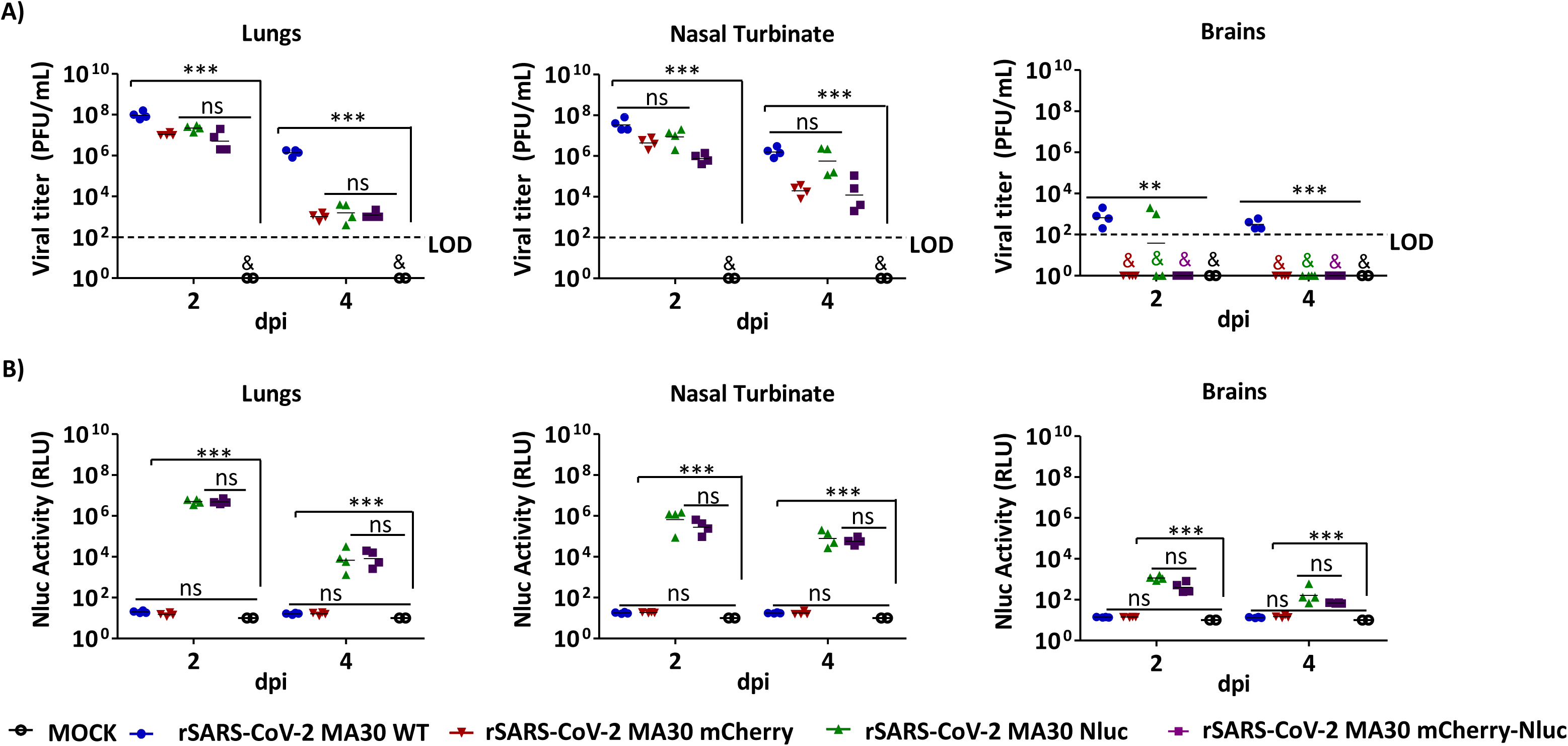
Viral titers and Nluc activity in tissue homogenates of C57BL/6 mice infected with reporter-expressing rSARS-CoV-2 MA30: Viral titers **(A)** and Nluc activity **(B)** in the lung (left), nasal turbinate (middle), and brain (right) tissue homogenates of 22-week-old C57BL/6 mice mock-infected or infected i.n. with 10^5^ PFU/mouse of rSARS-CoV-2 MA30 WT, rSARS-CoV-2 MA30 mCherry, rSARS-CoV-2 MA30 Nluc, and rSARS-CoV-2 MA30 mCherry-Nluc at 2 (n=4/group) and 4 (n=4/group) dpi were determined by standard plaque assay and microplate reader, respectively. The results are the mean values and SD. LOD, limit of detection. *\*\*p* < 0.01; *\*\*\*p* < 0.001; ns, not significant; (&) nd, not detected.

These results suggest that reporter-expressing rSARS-CoV-2 MA30, particularly those expressing Nluc, enabled sensitive detection of viral replication *in vivo* in intact animals and *ex vivo* in the lung and nasal turbinate homogenates of infected C57BL/6 mice. Likewise, rSARS-CoV-2 MA30 mCherry enabled detection of viral infection *ex vivo* in the lungs of infected animals at 2 dpi. However, viral titers of reporter-expressing rSARS-CoV-2 MA30 in lung, nasal turbinate and brain homogenates were lower than those observed in C57BL/6 mice infected with rSARS-CoV-2 MA30 WT, suggesting a slight attenuation of reporter-expressing rSARS-CoV-2 MA30 compared to rSARS-CoV-2 MA30 WT in C57BL/6 mice at both dpi due to expression of the foreign reporter genes.

### Characterization of reporter-expressing rSARS-CoV-2 MA30 in BALB/c mice

To further investigate the utility of the reporter-expressing rSARS-CoV-2 MA30 viruses in monitoring viral infection dynamics, we used the more susceptible BALB/c mouse model of infection (**Figure 3**). We infected 22-week-old female BALB/c mice i.n. with 10^5^ PFU of rSARS-CoV-2 MA30 WT, rSARS-CoV-2 MA30 mCherry, rSARS-CoV-2 MA30 Nluc, rSARS-CoV-2 MA30 mCherry-Nluc, or mock-infected (**Figure 7**). Nluc activity in mice at 1, 2, 4, and 6 dpi was evaluated by *in vivo* imaging (**Figure 7A**). Akin to our studies with C57BL/6 mice, we detected Nluc expression in BALB/c mice infected with the Nluc-expressing viruses (rSARS-CoV-2 MA30 Nluc and rSARS-CoV-2 MA30 mCherry-Nluc) (**Figure 7A**). Contrary to our previous results with C57BL/6 mice (**Figure 6A**), Nluc signals were observed as early as 1 dpi, especially in BALB/c mice infected with rSARS-CoV-2 MA30 mCherry-Nluc (**Figure 7A**). Nluc expression peaked at 2 dpi and began to decrease by 4 dpi. Notably, the Nluc signal was still detectable in BALB/c mice infected with rSARS-CoV-2 M30 mCherry-Nluc by 4 dpi, but it was lost in BALB/c mice infected with rSARS-CoV-2 MA30 Nluc (**Figure 7A**). Mice infected with rSARS-CoV-2 MA30 WT showed severe weight loss (**Figure 7B**), reaching approximately 75% of initial body weight by 7 dpi, similar to our previous studies (**Figure 3**). Interestingly, and parallel to the results in C57BL/6 mice, BALB/c mice infected with the reporter-expressing rSARS-CoV-2 MA30 exhibited less severe weight loss compared to C57BL/6 mice infected with rSARS-CoV-2 MA30 WT (**Figure 7B**). Comparable to the results in C57BL/6 mice, BALB/c mice infected with the rSARS-CoV-2 MA30 expressing reporter-genes loss ∼10% of their initial body weight but all of them recovered their body weight by 4-6 dpi, with all groups of mice reaching their initial body weight by 10 dpi (**Figure 7B**). Survival data (**Figure 7C**) indicated 100% mortality in BALB/c mice infected with rSARS-CoV-2 MA30 WT by 8 dpi, similar to our previous study (**Figure 3**). In contrast, all BALB/c mice infected with the reporter-expressing rSARS-CoV-2 MA30 survived the 10-day duration of the experiment, as did the mock-infected controls (**Figure 7C**). These results demonstrate that in BALB/c mice, resembling our previous studies in C57BL/6 mice, the reporter-expressing rSARS-CoV-2 MA30 is attenuated compared to rSARS-CoV-2 MA30 WT.

**Figure 7.**
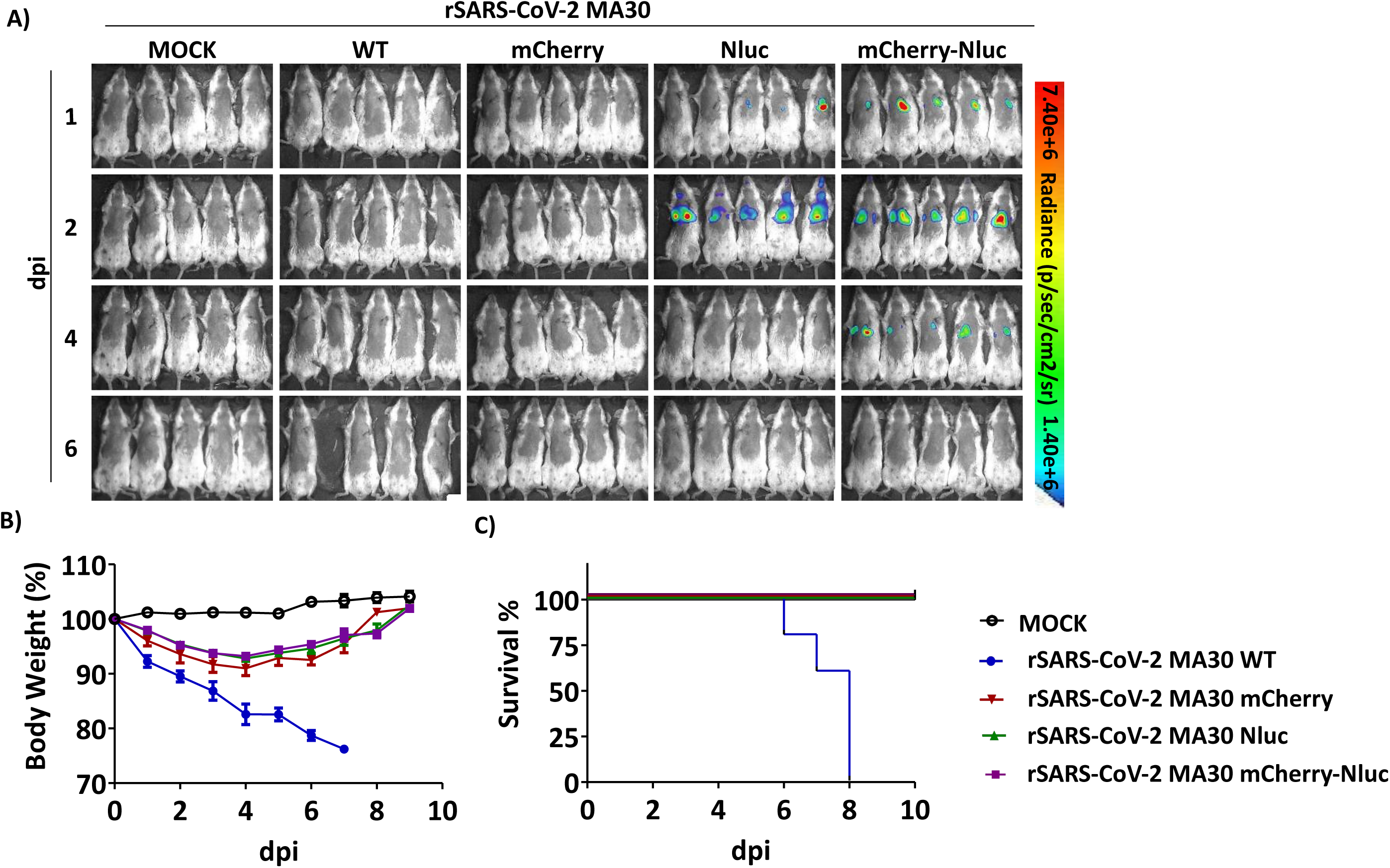
*In vivo* kinetics of reporter-expressing rSARS-CoV-2 MA30 in BALB/c mice: **A)** BALB/c 22-week-old mice (n=5/group) were mock-infected or infected i.n. with 10^5^ PFU/ mouse of rSARS-CoV-2 MA30 WT, rSARS-CoV-2 MA30 mCherry, rSARS-CoV-2 MA30 Nluc, or rSARS-CoV-2 MA30 mCherry-Nluc. Nluc expression in the whole mouse at the indicated dpi was evaluated with an Ami HT IVIS. Representative images of the same mouse at 1, 2, 4, and 6 dpi are shown. Changes in body weight **(B)** and survival **(C)** of mice infected in panel A were monitored for 10 days after viral infection.

*In vivo* and *ex vivo* detection of reporter rSARS-CoV-2 MA30 in BALB/c mice (**Figure 8**) revealed strong Nluc activity in mice infected with rSARS-CoV-2 MA30 Nluc and rSARS-CoV-2 MA30 mCherry-Nluc at 2 dpi, primarily localized in the lungs (**Figures 8A and 8B**). We were able to detect Nluc expression only in BALB/c mice infected with rSARS-CoV-2 MA30 mCherry-Nluc, but not rSARS-CoV-2 MA30 Nluc, at 4 dpi (**Figures 8C and 8D**). As expected, no Nluc signal was detected in mock-infected BALB/c mice or mice infected with rSARS-CoV-2 MA30 WT (**Figures 8A-8D**). *Ex vivo* imaging of excised lungs confirmed the presence of Nluc in lungs from mice infected with Nluc-expressing viruses, with signal intensity correlating with *in vivo* observations (**Figures 8B and 8D**): high levels of Nluc expression in the lungs of BALB/c mice infected with rSARS-CoV-2 MA30 Nluc and rSARS-CoV-2 MA30 mCherry-Nluc at 2 dpi (**Figure 8B**) and only Nluc signal in the lungs of BALB/c mice infected with rSARS-CoV-2 MA30 mCherry-Nluc, but not rSARS-CoV-2 MA30 Nluc, at 4 dpi (**Figure 8D**). Contrary to our previous results in C57BL/6 mice (**Figure 5**), we could not detect mCherry expression *ex vivo* in any of the lungs of BALB/c mice infected with rSARS-CoV-2 MA30 mCherry or rSARS-CoV-2 MA30 mCherry-Nluc at 2 or 4 dpi (**Figures 8B and 8D**).

**Figure 8.**
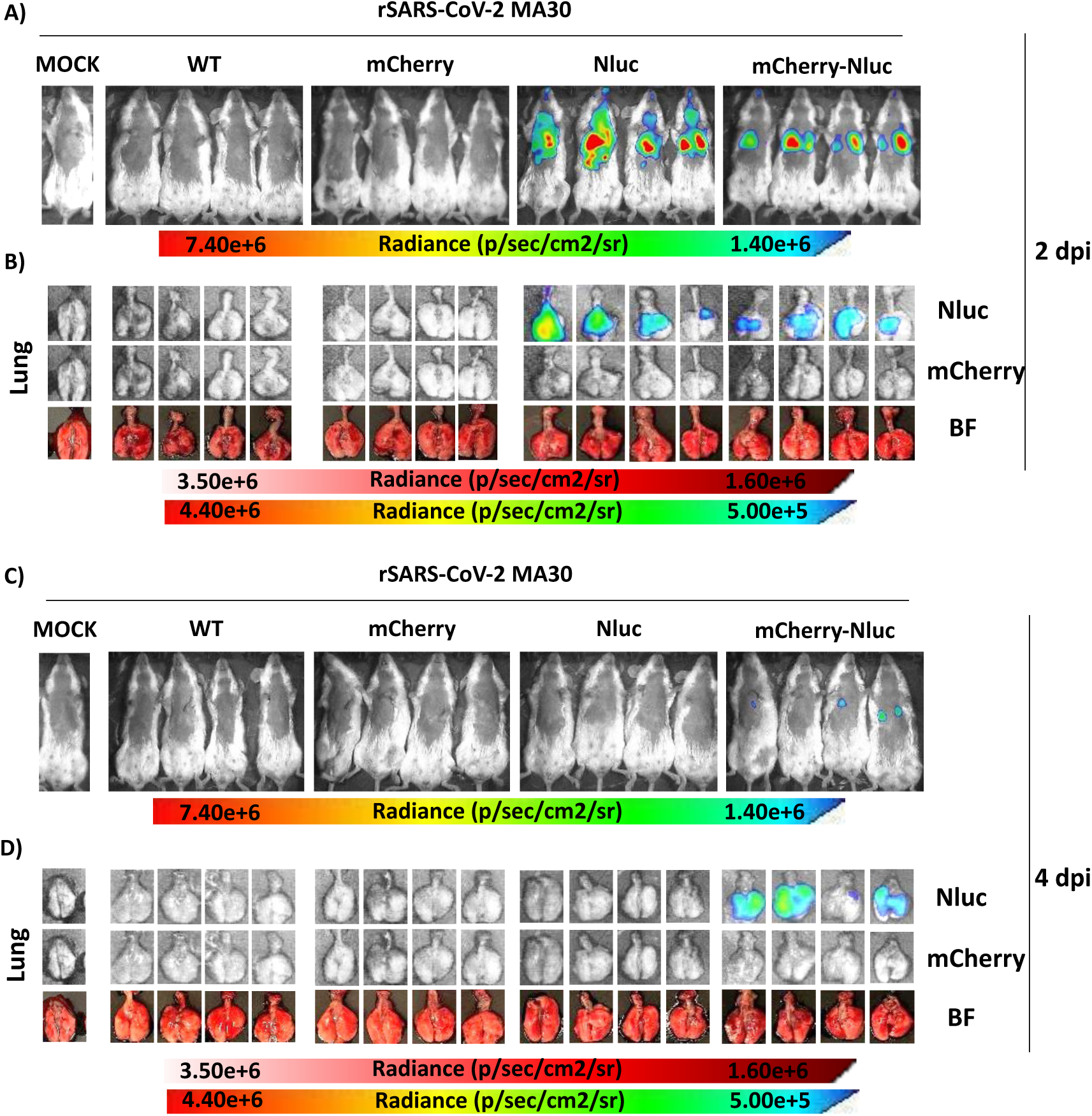
*In vivo* and *ex vivo* kinetics of reporter-expressing rSARS-CoV-2 MA30 in BALB/c mice: **A and C)** Nluc activity in whole 22-week-old BALB/c mice mock-infected or infected i.n. with 10^5^ PFU/mouse with rSARS-CoV-2 MA30 WT, rSARS-CoV-2 MA30 mCherry, rSARS-CoV-2 MA30 Nluc, and rSARS-CoV-2 MA30 mCherry-Nluc at 2 (n=4 mice/group) and 4 (n=4 mice/group) dpi using the Ami HT IVIS. **B and D)** Excised lungs from mock-infected or infected mice from panels A and C were monitored for Nluc and mCherry expression at 2 and 4 dpi. Bright field (BF) images of the lungs were also collected at the same time points.

Viral titers in tissue homogenates from the same infected BALB/c mice showed the highest levels of viral replication in the lungs of BALB/c mice infected with rSARS-CoV-2 MA30 WT at 2 dpi (∼10^7^-10^8^ PFU/mL), while reporter-expressing rSARS-CoV-2 MA30 showed slightly lower titers (∼10^7^ PFU/mL) by 2 dpi (**Figure 9A**). By 4 dpi, lung viral titers decreased significantly in all groups, with reporter viruses showing a more pronounced reduction (∼10^2^ PFU/mL) compared to rSARS-CoV-2 MA30 WT (∼10^5^ PFU/mL) (**Figure 9A**). Viral titers in the nasal turbinate followed a similar pattern, with rSARS-CoV-2 MA30 WT presenting higher titers at both 2 and 4 dpi compared to reporter viruses (**Figure 9A**). Akin to C57BL/6 mice (**Figure 6**), viral titers in the brain were low but detectable at 2 dpi in BALB/c mice infected with rSARS-CoV-2 MA30 WT and undetectable in all infected BALB/c mice by 4 dpi (**Figure 9A**). Quantification of Nluc activity in the same tissue homogenates showed the highest levels in the lungs and nasal turbinate of BALB/c mice infected with rSARS-CoV-2 MA30 Nluc and rSARS-CoV-2 MA30 mCherry-Nluc (**Figure 9B**). Lung Nluc activity peaked at 2 dpi and decreased by 4 dpi, while activity in nasal turbinate samples remained relatively stable at both dpi with both viruses Nluc (**Figure 9B**). No detectable Nluc activity was observed in brain homogenates at either time point (**Figure 9B**).

**Figure 9.**
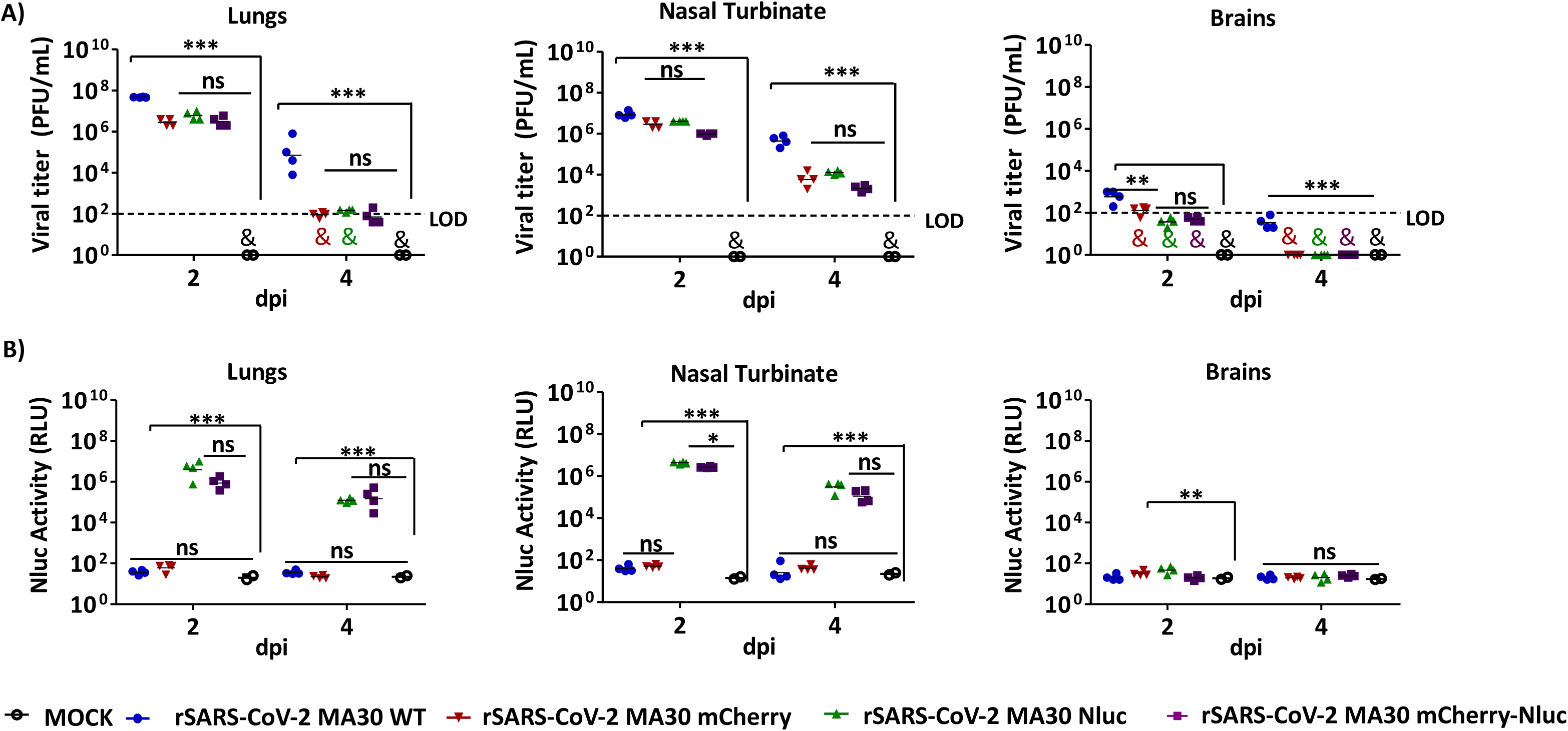
Viral titers and Nluc activity in tissue homogenates of BALB/c mice infected with reporter-expressing rSARS-CoV-2 MA30: Viral titers **(A)** and Nluc activity **(B)** in the lung (left), nasal turbinate (middle), and brain (right) tissue homogenates of 22-week-old C57BL/6 mice mock-infected or infected i.n. with 10^5^ PFU/mouse of rSARS-CoV-2 MA30 WT, rSARS-CoV-2 MA30 mCherry, rSARS-CoV-2 MA30 Nluc, and rSARS-CoV-2 MA30 mCherry-Nluc at 2 (n=4/group) and 4 (n=4/group) dpi were determined by standard plaque assay and a microplate reader, respectively. The results are the mean values and SD. LOD, limit of detection. *\*\*p* < 0.01; *\*\*\*p* < 0.001; ns, not significant; (&) nd, not detected.

These results demonstrate that reporter-expressing rSARS-CoV-2 MA30 enables detection and quantification of viral replication *in vivo* and *ex vivo* in BALB/c infected mice. However, and parallel to the results in C57BL/6 mice, the reporter-expressing rSARS-CoV-2 MA30 are attenuated in BALB/c mice compared to rSARS-CoV-2 MA30 WT.

## Discussion

The development and characterization of reporter-expressing rSARS-CoV-2 MA30 in this study opens new possibilities for preclinical studies with SARS-CoV-2. These reporter viruses, which express mCherry and Nluc, alone or together, facilitate real-time *in vitro* as well as *in vivo*, or *ex vivo*, tracking of viral replication and infection dynamics in cultured cells and C57BL/6 and BALB/c mouse models, offering new opportunities to evaluate prophylactic and/or therapeutic interventions without the need for secondary approaches to detect the presence of the virus or the use of K18-hACE2 transgenic mice.

Our results suggest that cloning or expression of mCherry, Nluc, and mCherry-Nluc from the locus of the N protein resulted in slight viral attenuation compared to rSARS-CoV-2 MA30 WT. This attenuation was evidenced by slightly reduced viral plaque sizes in Vero AT and Vero E6 cells (**Figure 1**), lower peak viral titers in Vero AT and A549 hACE2 (**Figure 2**), and reduced weight loss and mortality *in vivo* compared to rSARS-CoV-2 MA30 WT in both C57BL/6 (**Figure 4**) and BALB/c (**Figure 7**) mice. Importantly, attenuation did not compromise reporter gene expression *in vitro* (**Figure 2**), *in vivo* (**Figures 4 and 7**), or *ex vivo* in C57BL/6 and BALB/c (**Figures 5 and 8**) mice infected with the reporter-expressing rSARS-CoV-2 MA30. Similar findings have been reported for other reporter-expressing viruses, such as vesicular stomatitis virus (VSV) and respiratory syncytial virus (RSV), where insertion and/or expression of reporter genes affected viral fitness and pathogenicity *in vivo* (35, 36). This contrasts with our previous studies where cloning and expression of the same or similar reporter genes, alone or in combination, using the same PTV-1 2A approach, from the locus of the N protein, did not result in attenuation of SARS-CoV-2 WA1 *in vivo* in K18 hACE2 mice (28, 29, 37). This can be a result of the high susceptibility of K18 hACE2 mice to infection with SARS-CoV-2 WA1 (38). While attenuation is not a desirable characteristic for recombinant viruses expressing reporter genes, it opens the feasibility of still being able to track viral infection *in vitro*, *ex vivo*, or *in vivo*, without exceeding virulence. Actually, the lack of excessive virulence of the reporter-expressing rSARS-CoV-2 MA30 compared to rSARS-CoV-2 MA30 WT in both C57BL/6 and BALB/c mice (**Figure 3**) could represent an advantage in studies aimed at determining long COVID or in studies to identify host factors associated with viral attenuation by using WT and knockout BALB/c or C57BL/6 mice.

The robust expression of mCherry and Nluc reporters in infected cells and tissues underscores the utility of these viruses for real-time tracking of SARS-CoV-2 infection. IVIS revealed dynamic changes in viral replication, with peak signals at 2 dpi in both C57BL/6 and BALB/c mice that correlate with the peak of viral titers. Reporter gene expression *in vitro*, *ex vivo*, or *in vivo* offers a significant advantage over traditional methods to detect the presence of the virus. Notably, non-invasive Nluc expression using IVIS provides an unparalleled advantage for studying the spatiotemporal dynamics of SARS-CoV-2 infection in WT C57BL/6 and BALB/c mice. We and others have previously used this approach on other viral models, including influenza viruses, to track and monitor viral dissemination in the entire mouse (39, 40). Importantly, Nluc expression correlated well with viral titers in lungs and nasal turbinate of infected mice, validating its application for quantitative studies of SARS-CoV-2 dynamics. And analogous to our results with reporter-expressing rSARS-CoV-2 MA30 in this study, expression of reporter genes from influenza viruses resulted in attenuation compared to their WT counterparts (41).

Reporter-expressing rSARS-CoV-2 MA30 also has a significant potential for evaluating prophylactics and therapeutics, where reporter gene expression (Nluc) provides a sensitive and quantitative rapid assessment of protection and antiviral activity *in vivo*, allowing for the interrogation of prophylactic vaccines or therapeutic antivirals using a reduced number of animals. Additionally, the ability to visualize viral dynamics in live animals using Nluc expression and IVIS provides insights into therapeutic timing, dosing, and efficacy, as well as longitudinally evaluating viral replication in the same animals, facilitating the optimization and/or comparison of intervention strategies.

An intriguing observation was the robust detection of viral titers in brain tissue homogenates at 2 dpi in rSARS-CoV-2 MA30 WT-infected mice (∼10^3^ and ∼10^2^-10^3^ PFU in C57BL/6 and BALB/c, respectively), despite the absence of neuropathology or clinical neurological signs. Reporter-expressing rSARS-CoV-2 MA30 Nluc showed reduced brain tropism (2 of 4 C57BL/6 mice with detectable virus by 2 dpi), paralleling their attenuated pulmonary phenotype. Notably, *ex vivo* imaging revealed no detectable mCherry or Nluc signal in brain tissue despite robust reporter expression in lungs, suggesting that infectious virus in the brain is sequestered in cerebral vasculature rather than replicating in parenchymal cells (42). This vascular sequestration hypothesis is supported by: (i) rapid early brain detection (2 dpi) without pathology; (ii) reduced brain tropism in attenuated reporter viruses; and (iii) rapid viral clearance by 4 dpi. Future studies using immunofluorescence with vascular markers (CD31), intravital two-photon microscopy, or brain vasculature microdissection would directly test this hypothesis (43). These mechanistic investigations represent a valuable avenue for future work but are beyond the scope of the current characterization study.

While the reporter-expressing rSARS-CoV-2 MA30 described in this study offers numerous advantages over rSARS-CoV-2 MA30 WT, they also present limitations. The insertion and/or expression of reporter genes, while resulting in minimal attenuation *in vitro*, significantly affects viral pathogenicity and replication *in vivo* compared to rSARS-CoV-2 MA30 WT. Future studies aimed to determine the molecular mechanism of *in vivo* attenuation are guaranteed to provide reporter-expressing rSARS-CoV-2 MA30 with an *in vivo* phenotype resembling that of rSARS-CoV-2 MA30 WT. Another limitation is the stability of reporter gene expression during serial passaging. Although our study confirmed the integrity of reporter genes *in vitro* and *in vivo*, stability studies to assess the stability of the reporter gene after serial passages in cultured cells should be evaluated to ensure reporter gene expression after viral passages. Previous studies using reporter-expressing rSARS-CoV-2 WA1 with the same PTV-1 2A approach from the locus of the viral N protein supported the genetic stability of this strategy. Finally, while our study focused on mouse models, the application of these reporter-expressing rSARS-CoV-2 MA30 in other animal models, such as hamsters (44, 45), dwarf hamsters (46), or ferrets (47, 48), could provide additional insights into the feasibility of their use in other animal models of SARS-CoV-2 infection.

Altogether, the reporter-expressing rSARS-CoV-2 MA30 developed in this study expands the research tools for studying SARS-CoV-2. Their ability to provide real-time, quantitative measurements of viral replication without the need for secondary approaches to detect the presence of the virus in infected cells or BALB/c and C57BL/6 mice opens the feasibility of extending SARS-CoV-2 viral replication, host-pathogen interactions, and the feasibility of evaluating new vaccines and/or antivirals for the prevention and treatment of SARS-CoV-2 infection.

## Materials and Methods

### Biosafety and ethics statement

*In vitro* and *in vivo* experiments involving infectious parental or recombinant SARS-CoV-2 were conducted in biosafety level 3 (BSL3) and animal BSL3 (ABSL3) laboratories at Texas Biomedical Research Institute. The Texas Biomedical Research Institute Biosafety and Recombinant DNA Committees (BSC and RDC, respectively) and the Institutional Animal Care and Use Committee (IACUC; protocol no. 1718MU) approved the experimental procedures involving cell culture and animal studies, respectively. Mice were maintained in the animal care ABSL3 facility at Texas Biomedical Research Institute under specific pathogen-free conditions.

### Cells

African green monkey kidney epithelial Vero E6 cells (CRL-1586), were obtained from the American Type Culture Collection (ATCC; Bethesda, MD). Human A549 cells expressing hACE2 (A549 hACE2) and Vero E6 cells expressing hACE2 and TMPRSS2 (Vero AT) were obtained from BEI Resources (NR-53821 and NR-54970, respectively). Cells were maintained in Dulbecco’s modified Eagle medium (DMEM; Corning) supplemented with 5% (vol/vol) fetal bovine serum (FBS; VWR) and 1% PSG (100 U/mL penicillin, 100 μg/mL streptomycin, and 2 mM l-glutamine; Corning) at 37°C in a 5% CO_2_ incubator. Vero AT cells were treated with 5 μg/mL puromycin every other passage. A549 hACE2 cells were treated with 5 μg/mL of Plasticidin every other passage.

### Generation of WT and reporter-expressing rSARS-CoV-2 MA30

The rSARS-CoV-2 MA30 WT and expressing reporter genes were generated using the previously described bacterial artificial chromosome (BAC)-based reverse genetics system (11, 28, 37, 49–51) of the MA30 strain of SARS-CoV-2 Washington-1 isolate, WA1 (USA-WA1/2020) (21). SARS-CoV-2 MA30 was generated by serial passaging WA1 in BALB/c mice (21). The SARS-CoV-2 MA30 BAC WT backbone was used to generate reporter viruses expressing mCherry (rSARS-CoV-2 MA30 mCherry), nanoluciferase (rSARS-CoV-2 MA30 Nluc), and a fusion of mCherry-Nluc (rSARS-CoV-2 MA30 mCherry-Nluc) from the locus of the viral N protein as previously described (9, 17, 28). All reporter-expressing rSARS-CoV-2 MA30 generated in this study are based exclusively on the MA30 backbone; no hybrid viruses containing mutations from both SARS-CoV-2 MA10 and SARS-CoV-2 MA30 were constructed. The reporter genes were inserted into the backbone of the SARS-CoV-2 MA30 genome, ensuring that any observed changes in viral fitness or pathogenicity are attributable to the insertion of reporter genes. BAC plasmids were used for viral rescue in Vero E6 cells as previously described (16). Briefly, Vero E6 cells (1.2 × 10^6^ cells/well, 6-well plate format) were transfected with 4.0 μg/well of the BAC using Lipofectamine 2000 (Thermo Fisher Scientific). After overnight transfection, the transfection media were changed to post-infection media (DMEM containing 2% FBS and 1% PSG), and cells were scaled up into T75 flasks 2 days post-transfection. After 3 days, rescue of reporter-expressing rSARS-CoV-2 MA30 was evaluated by the presence of cytopathic effect (CPE), fluorescence (rSARS-CoV-2 MA30 mCherry and rSARS-CoV-2 MA30 mCherry-Nluc), and Nluc (rSARS-CoV-2 MA30 Nluc and rSARS-CoV-2 MA30 mCherry-Nluc) expression. Cell culture supernatants were collected, labeled as P0, and stored at −80°C. After viral titration by plaque assay, P1 viral stocks were generated by infecting fresh Vero E6 cells at a low multiplicity of infection (MOI; 0.0001 PFU/cell) for 3 days and stored at −80°C. Viral titers (PFU/mL) were determined by plaque assay in Vero E6 and Vero AT cells.

### Plaque assays

Vero E6 and Vero AT cells (1.2 × 10^6^ cells/well, 6-well format) were infected with rSARS-CoV-2 MA30 WT, rSARS-CoV-2 MA30 mCherry, rSARS-CoV-2 MA30 Nluc, or rSARS-CoV-2 MA30 mCherry-Nluc. After 1 hour of viral absorption, viruses were removed, and cells were overlaid with post-infection media containing 0.6% agar (Oxoid) and incubated at 37°C in a 5% CO_2_ incubator. At 72 hours post-infection (hpi), cells were fixed overnight at 4°C in 10% neutral buffered formalin. For detection of mCherry expression, fixed cells were imaged using a ChemiDoc MP imaging system (Bio-Rad) to detect mCherry-positive plaques. For detection of Nluc, fixed cells were incubated with PBS containing the Nano-Glo luciferase substrate following the manufacturer’s instructions (Promega Nano-Glo Luciferase Assay System). Nluc-expressing plaques were visualized using a ChemiDoc MP imaging system (Bio-Rad). Following detection of mCherry and Nluc signals, cells were permeabilized with T-PBS (0.05% Tween-20 in PBS) for 10 minutes at room temperature. Nonspecific binding sites were blocked using 2.5% bovine serum albumin (BSA) in PBS for 1 hour. Cells were then incubated with a primary mouse monoclonal antibody (MAb) against SARS-CoV N protein (1C7C7) (17) or with a rabbit polyclonal (PAb, Promega) against Nluc for 1 hour at room temperature. Secondary staining was performed using anti-mouse and anti-rabbit Vectastain ABC kits, followed by visualization with the DAB HRP Substrate according to the manufacturer’s instructions (Vector Laboratories). Stained plaques were imaged using a ChemiDoc MP imaging system (Bio-Rad). For Vero-AT cells, plaques were visualized using crystal violet staining. After cell fixation, cells were washed twice with PBS and stained with a crystal violet solution (0.5% crystal violet in 20% ethanol) for 30 minutes at room temperature. Next, plates were gently washed with tap water to remove the excess of crystal violet staining, and viral plaques were visualized as clear zones against the stained monolayer. Plaques visualized using fluorescence, chemiluminescence, immunostaining, or crystal violet were manually quantified.

### Viral growth kinetics

Vero AT and A549 hACE2 cells (0.5 × 10^6^ cells/well, 12-well plate format, triplicates) were infected (MOI, 0.01) with rSARS-CoV-2 MA30 WT, rSARS-CoV-2 MA30 mCherry, rSARS-CoV-2 MA30 Nluc, and rSARS-CoV-2 MA30 mCherry-Nluc. After 1 hour of viral adsorption, cells were washed with PBS and incubated at 37°C in post-infection media. Viral titers in cell culture supernatants at each indicated time point (12, 24, 48, and 72 hpi) were determined by plaque assay as described above. At each time point, mCherry expression was visualized with an EVOS M5000 imaging system. Nluc activity in the cell culture supernatants at the same times post-infection was quantified using a microplate reader and a Nano-Glo Luciferase Assay system following the manufacturer’s recommendations (Promega). Mean values and standard deviation (SD) were calculated with GraphPad Prism.

### Mouse lethal dose 50 (MLD_50_)

To determine age-dependent and strain-specific susceptibility to rSARS-CoV-2 MA30 across different mouse models, 6- and 21-week-old females C57BL/6, BALB/c, and K18-hACE2 mice (n=4/group) (The Jackson Laboratory) were anesthetized using isoflurane and subsequently infected intranasally (i.n.) with 10-fold serial dilutions (10²–10 PFU/mouse in 50 µL PBS) of rSARS-CoV-2 MA30. Mice were monitored daily for 10 days for body weight changes, clinical signs (hunching, ruffled fur, labored breathing), and survival. Mice that were below 75% of their initial body weight were considered to have reached their experimental endpoint and were humanely euthanized. The median lethal dose (MLD_50_) was calculated using the Reed-Muench method (52).

### *In vivo* and *ex vivo* analysis of infection dynamics

Female 20- to 22-week-old C57BL/6 and BALB/c mice (The Jackson Laboratory) were anesthetized intraperitoneally (i.p.) with a mixture of Ketamine (100 mg/mL) and Xylazine (20 mg/mL). For body weight and survival studies, C57BL/6 or BALB/c mice (n = 5/group) were infected i.n. with 10^5^ PFU/mouse of the indicated viruses and monitored daily for body weight loss and survival to assess morbidity and mortality, respectively, for 10 days. Mice that were below 75% of their initial body weight were considered to have reached their experimental endpoint and were humanely euthanized. A separate group of mice was also mock-infected with PBS and served as a negative control. Survival data were analyzed using Kaplan-Meier curves (53). *In vivo* Nluc imaging of live mice was conducted with an Ami HT *in vivo* imaging system (IVIS; Spectral Instruments) at 1-, 2-, 4-, and 6-days post-infection (dpi). At each time point, mice were anesthetized i.p. with a mixture of Ketamine (100 mg/mL) and Xylazine (20 mg/mL) and retro-orbitally injected with 100 μL of Nano-Glo luciferase substrate (Promega) diluted 1:10 in PBS. Mice were immediately placed in an isolation chamber and imaged using the Ami HT IVIS. Bioluminescence data were acquired and analyzed using the Aura software (AMI Spectral Instruments), and total flux values (protons/s) were normalized to the background signal of the mock-infected control. To determine viral titers and assess fluorescence expression in the lungs, a separate cohort of mice (n = 8/group) was similarly infected with the indicated recombinant viruses and humanely euthanized at 2 (n = 4/group) and 4 (n = 4/group) dpi after *in vivo* imaging with the Ami HT IVIS. Lungs were surgically excised, washed in PBS, and bioluminescence and fluorescent images were obtained using the Ami HT IVIS. Lung brightfield images were obtained using an iPhone. Nasal turbinate (NT), lung, and brain tissues were individually homogenized in 1 mL of PBS using a Precellys tissue homogenizer (Bertin Instruments) for 20 seconds at 7,000 rpm. Tissue homogenates were then centrifuged at 12,000 × g at 4°C for 5 minutes to pellet cell debris, and supernatants were collected. Viral titers were determined by plaque assay and immunostaining, as described above. Nluc activity in the tissue homogenates was measured using the Nano-Glo luciferase substrate kit (Promega) and a Glowmax microplate reader.

### Statistical analysis

All data are presented as mean values and SD for each group and were analyzed using GraphPad Prism software version 9.5.1 (GraphPad Software, LLC, USA). A one-way analysis of variance (ANOVA) and Tukey’s test for multiple-comparison correction were used for statistical analysis on GraphPad Prism. Statistical significance was as follows: *, *P* < 0.05; **, *P* < 0.01; ***, *P* < 0.001; ****, *P* < 0.0001; and ns, no significance.

## Acknowledgments

We thank BEI Resources for providing Vero AT (NR-54970) and A549 hACE2 (NR-53821) cells.

